# Discovery and engineering of small SlugCas9 with broad targeting range and high specificity and activity

**DOI:** 10.1101/2020.09.29.316661

**Authors:** Ziying Hu, Chengdong Zhang, Shuai Wang, Jingjing Wei, Miaomiao Li, Linghui Hou, Hongmao Liu, Dong Liu, Feng Lan, Daru Lu, Hongyan Wang, Jixi Li, Yongming Wang

## Abstract

The compact CRISPR/Cas9 system, which can be delivered by adeno-associated virus (AAV), is a promising platform for therapeutic applications. However, current compact Cas9 nucleases have limited activity, targeting scope and specificity. Here, we identified three compact SaCas9 orthologs, *Staphylococcus lugdunensis* Cas9 (SlugCas9), *Staphylococcus lutrae* Cas9 (SlutrCas9) and *Staphylococcus haemolyticus* Cas9 (ShaCas9), for mammalian genome editing. Interestingly, SlugCas9 recognizes a simple NNGG PAM and displays comparable activity to SaCas9. We further generated a SlugCas9-SaCas9 chimeric nuclease, which has both high specificity and high activity. We lastly engineered SlugCas9 with mutations to generate a high fidelity variant that maintains high specificity without compromising on-target editing efficiency. Our study offers important minimal Cas9 tools that are ideal for both basic research and clinical applications.

## Introduction

Compact CRISPR/Cas9 nucleases (<1,100 aa) can be delivered by adeno-associated virus (AAV) ^1-4^ for *in vivo* genome editing, and thus hold great promise for gene therapy. The active complex is composed of a Cas9 nuclease and a guide RNA (gRNA), which together recognize a target DNA that is complementary to the 20 bp-protospacer sequence in the gRNA. Upon recognition, this complex generates a site-specific double-strand break (DSB) ^5-10^. Target site recognition also requires a specific protospacer adjacent motif (PAM) ^5, 11-13^ unique to the Cas9 protein, which limits the targeting scope of Cas9. We and others have identified several compact Cas9 nucleases for mammalian genome editing ^2, 14-18^. However, none maintains the combination of desired properties such as broadened PAM targeting range, high activity and high specificity.

In this study, we identified three compact SaCas9 orthologs–*Staphylococcus lugdunensis* Cas9 (SlugCas9), *Staphylococcus lutrae* Cas9 (SlutrCas9) and *Staphylococcus haemolyticus* (ShaCas9), for mammalian genome editing. Initial SaCas9 proteins require a long and complex PAM for targeting (NNGRRT for SaCas9, NNNRRT for SaCas9-KKH) ^2, 19^, which limits their widespread usage. These new *Staphylococcus* Cas9 proteins recognize an NNGG, NNGRR and NNGGV (V=A or C or G) PAM, respectively. The most interesting SlugCas9 contains a compact PAM recognition motif, analogous to the NGG PAM of the commonly used SpCas9, and is ∼1kp smaller in size. To reduce the propensity of off-target cleavage, we generated a SlugCas9-based chimeric Cas9 nuclease with high specificity and activity. Importantly, we also engineered a variant of SlugCas9 to be of high-fidelity, hereby referred to as SlugCas9-HF. This new Cas protein encompasses all desired properties such as recognizing an NNGG PAM, displays high specificity and maintains high activity.

## Results

### Identification of SaCas9 orthologs for genome editing

To identify novel compact Cas9 nucleases for genome editing, we used *Staphylococcus aureus* Cas9 (SaCas9) as a reference and searched for related orthologs from UniProt ^20^. We focused on Cas9 nucleases with relatively high identity (at least 50%) to SaCas9 and subsequently selected five orthologs for characterization, including Staphylococcus equorum Cas9 (SeqCas9, 97.1% identity), Staphylococcus lugdunensis Cas9 (SlugCas9, 63.2% identity), Staphylococcus epidermidis Cas9 (SepCas9, 64.2% identity), Staphylococcus haemolyticus Cas9 (ShaCas9, 63.2% identity) and Staphylococcus lutrae Cas9 (SlutrCas9, 59.1% identity). Importantly, these Cas9 orthologs differ in at least one residue located at the protein domain that is important for PAM recognition, corresponding to residues N986 and R991 in SaCas9 (Figure 1A) ^21^, suggesting that these orthologs may recognize different PAM motifs.

**Figure 1.**
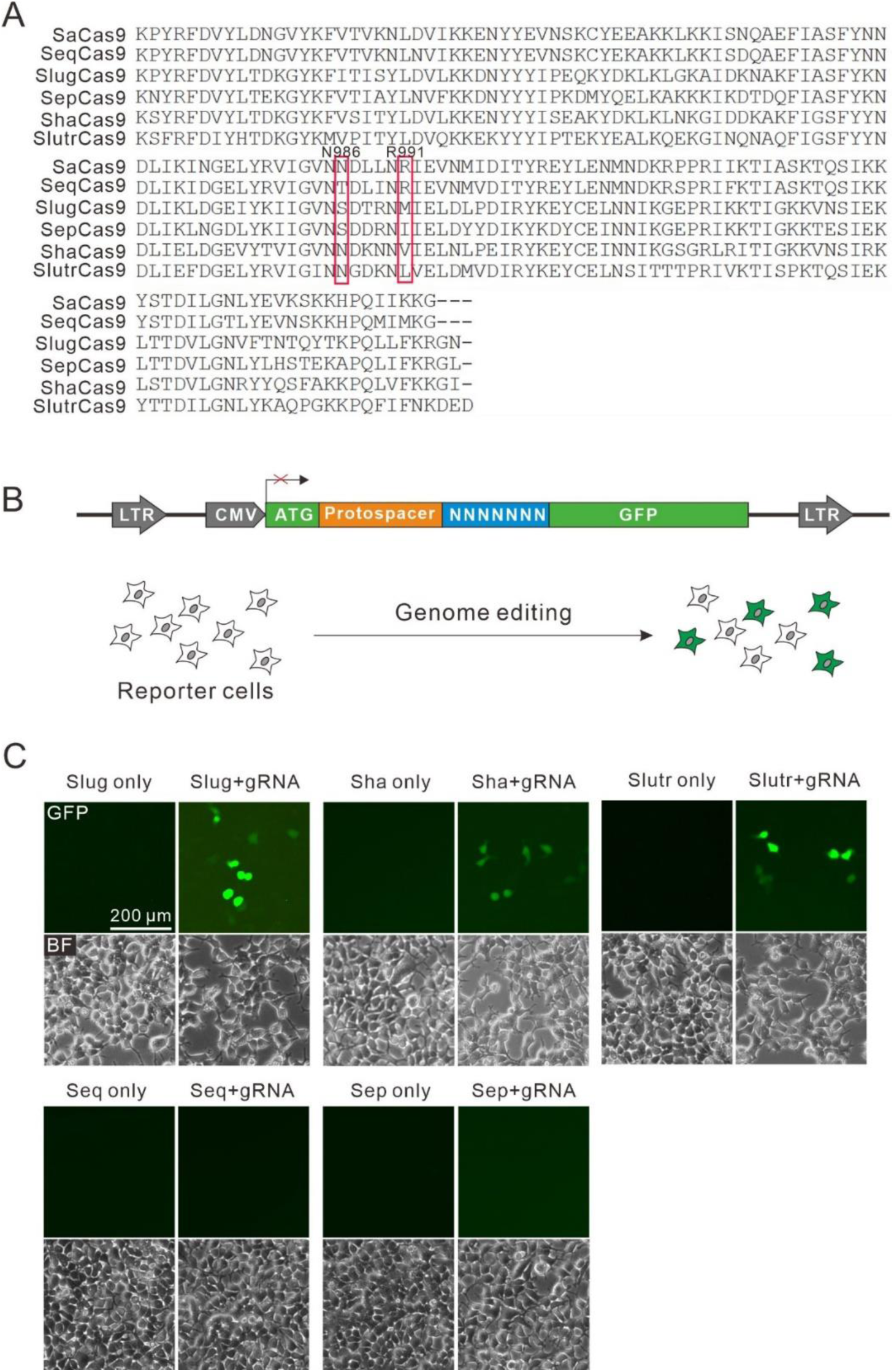
A GFP-activation assay reveals that SlugCas9, ShaCas9 and SlutrCas9 enable genome editing. (A) PAM interaction domain (PID) sequence alignment of SaCas9 orthologs. Amino acids important for PAM recognition are indicated by red box. (B) Schematic of the GFP-activation assay for Cas9 activity testing. A GFP reporter is disrupted by a protospacer followed by a 7-bp random sequence between ATG and GFP coding sequence. The reporter library is stably integrated into HEK293T cells. Genome editing will induce GFP expression for a portion of cells. (C) Transfection of SlugCas9, ShaCas9 and SlutrCas9 with gRNAs induce GFP expression.

We first used a previously reported GFP-activation PAM screening assay (Figure 1B) ^18^ to evaluate active genome editing and to identify the Cas9s’ respective PAMs. Each Cas9 ortholog was human-codon optimized, synthesized and cloned into a mammalian SaCas9 expression plasmid construct. The canonical SaCas9 gRNA scaffold was used for gRNA expression ^2^. Transfection of SlugCas9, ShaCas9 or SlutrCas9 with gRNAs could induce GFP expression (Figure 1C), suggesting that they indeed enabled active genome editing in mammalian cells. GFP-positive cells were subsequently isolated by flow cytometry and the target DNA was PCR-amplified for deep-sequencing.

Sequencing results revealed that insertions and deletions (indels) occurred at the target sites for these three Cas9 nucleases at sites encoded in their respective protospacers (Figure 2A). WebLogo revealed that SlugCas9 preferred G-rich PAM (Figure 2B), and a PAM wheel revealed that SlugCas9 preferred an NNGG PAM targeting motif (Figure 2C). Both WebLogo and the PAM wheel revealed that ShaCas9 and SlutrCas9 preferred an NNGGV (V=A or C or G) and NNGRR (R=A or G) PAM, respectively (Figure 2D-G).

**Figure 2.**
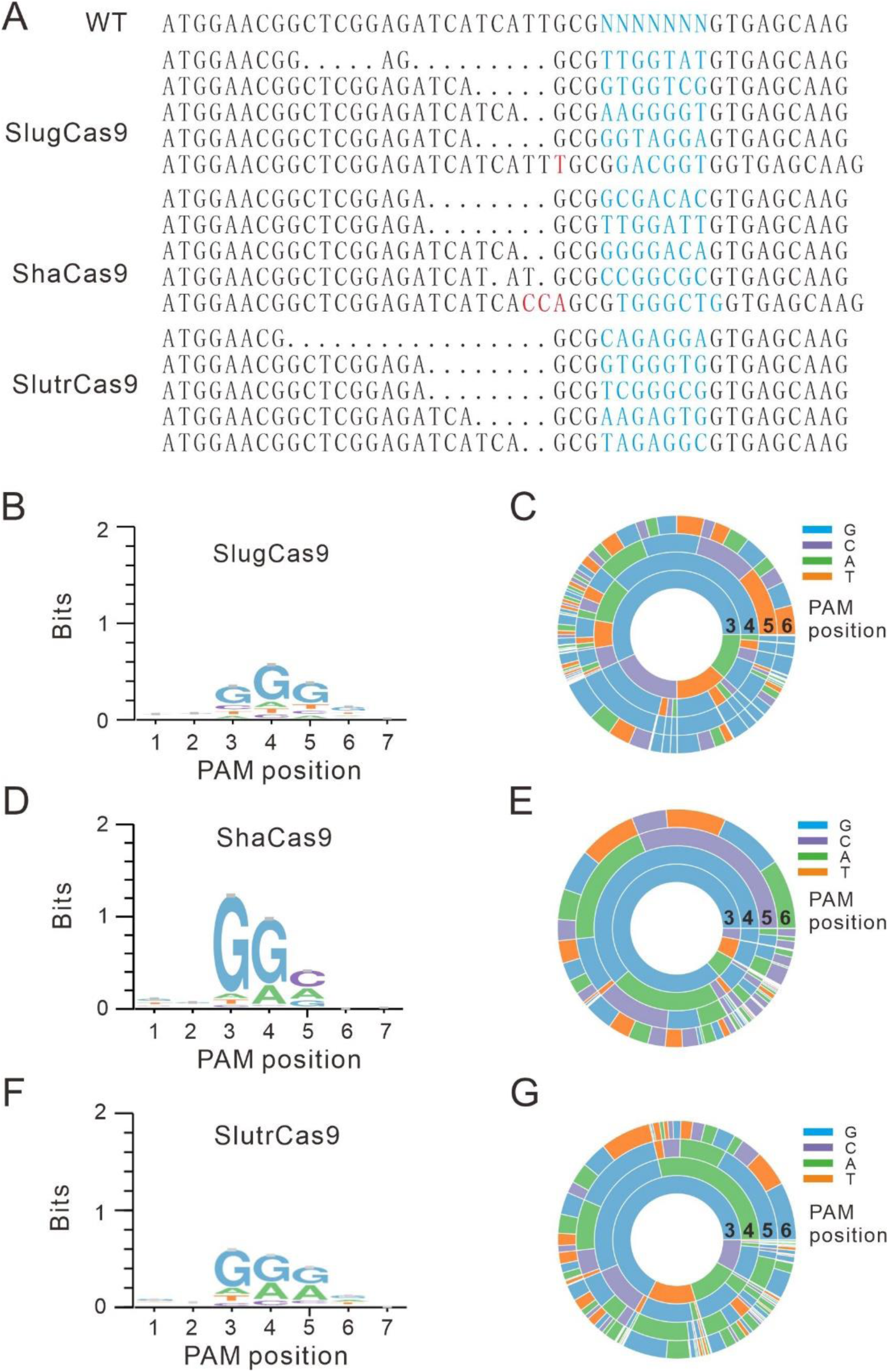
PAM sequence analysis. (A) Deep sequencing reveals that SlugCas9, ShaCas9 and SlutrCas9 generate indels on the targets. (B, D, F) WebLogos for SlugCas9, ShaCas9 and SlutrCas9 are generated based on deep sequencing data. (C, E, G) PAM wheels for SlugCas9, ShaCas9 and SlutrCas9 are generated based on deep sequencing data.

### SaCas9 orthologs enable genome editing for endogenous loci

To test the genome editing capability of these three Cas9 nucleases, we selected a panel of endogenous targets loci. All three Cas9 nucleases efficiently generated indels in HEK293T cells (Figure 3A) at the respective sites. We focused further analysis on SlugCas9 due to its short and simple PAM recognition motif. Selected target sites in Figure 3A contained an NNGRRT PAM that can be edited by SaCas9, allowing for a side-by-side comparison of editing efficiency at the same site. We used the SaCas9 plasmid backbone for the three Cas9 orthologs, and confirmed by qRT-PCR to note similar expression levels of SlugCas9 and SaCas9. The editing results revealed that SlugCas9 and SaCas9 displayed similar efficiency at 14 endogenous loci (Figure 3A and 3B).

**Figure 3.**
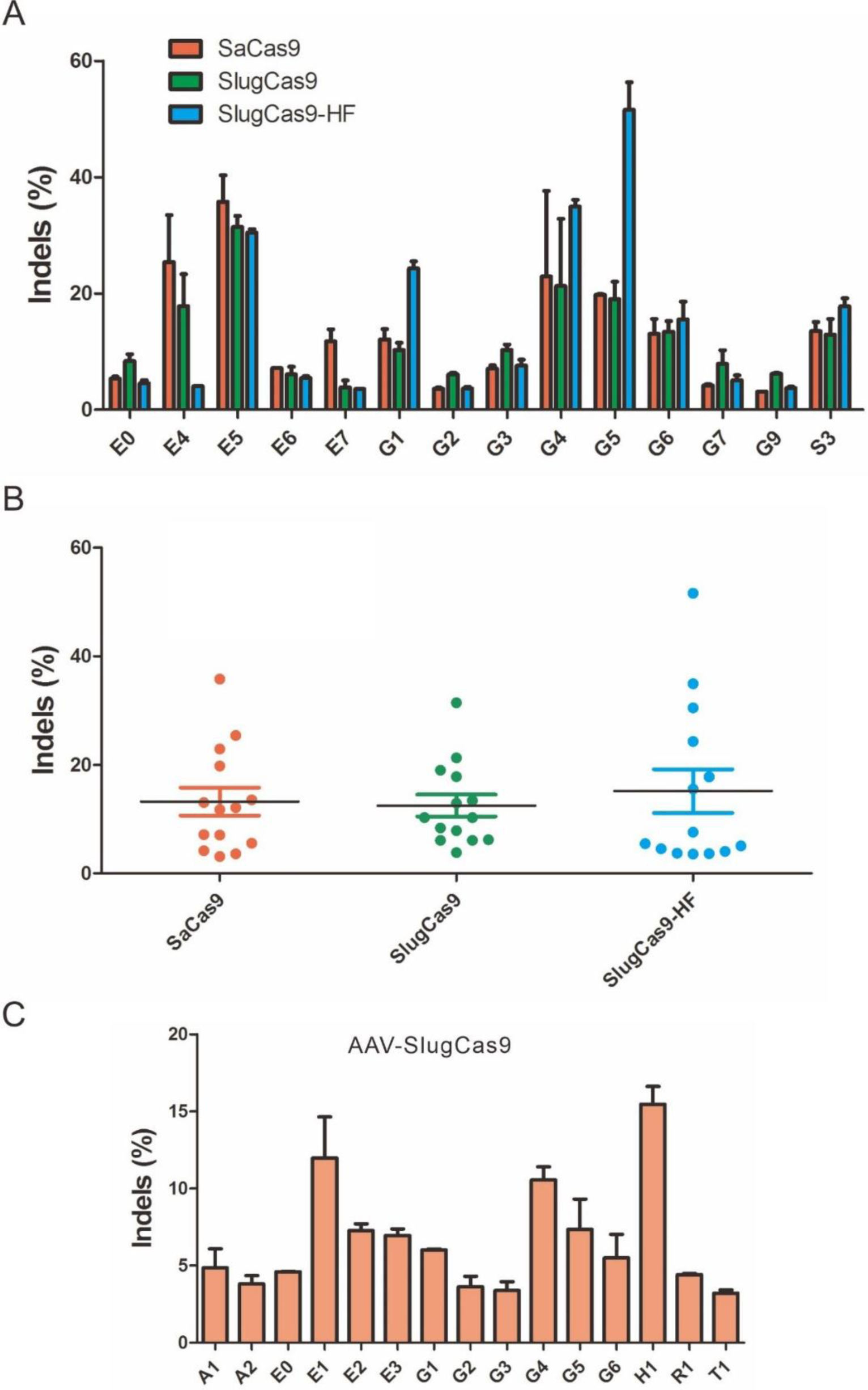
Genome editing for endogenous sites. (A) Comparison of SaCas9, SlugCas9 and SlugCas9-HF efficiency for genome editing at 14 endogenous loci. n=2. (B) Quantification of editing efficiency for SaCas9, SlugCas9 and SlugCas9-HF. (C) SlugCas9 can be delivered by AAV for genome editing in HEK293T cells. n=2.

We further tested the genome editing capacity of SlugCas9 in additional cell types, including A375, HCT116, HeLa, human foreskin fibroblast (HFF, primary cells) cells and N2a (mouse neuroblastoma cell line) cells. SlugCas9 generated indels in all these cell types with varying efficacies. To facilitate delivery efforts, we packaged SaCas9 together with its gRNA into AAV and infected HEK293T, A375, HCT116, HeLa and HFF cells. Indels were detected in all cell types, but frequencies varied depending on the loci and cell types (Figure 3C). We further identified the SlugCas9 trans-activating CRISPR RNA (tracrRNA) and established a single guide RNA (sgRNA) for genome editing. Since there were four continuous nucleotides of Ts in the sgRNA, which could lead to premature termination ^22^, we mutated the fourth T to C to increase sgRNA expression. SlugCas9 was transfected together with either wild-type or engineered gRNA into HEK293T cells for genome editing. Although both gRNAs enabled editing, the engineered gRNAs were more efficient at three out of four tested loci. Taken together, we recommend SlugCas9 as a minimal and efficient enzyme for genome editing.

### Base editing with SlugCas9

Base editing is a powerful technology that enables the programmable conversion of an A:T base pair to G:C or a C:G base pair to T:A in the mammalian genome ^23-26^. This technology utilizes the fusion of a catalytically disabled Cas9 nuclease to a nucleobase deaminase enzyme. To decrease protein size and facilitate delivery efforts, SaCas9, which is ∼1 kb smaller than SpCas9, has been used for base editing ^27^. To test whether SlugCas9 can be employed for base editing, we generated a nickase form of SlugCas9 (SlugCas9n) by introducing a D10A mutation. We replaced the nickase form of SpCas9 with SlugCas9n in BE4max ^28^ to generate APOBEC1–SlugCas9n–UGI (SlugBE4max, Figure 4A). We tested the editing capability of SlugBE4max at a panel of 34 endogenous sites. SlugBE4max and the respective gRNAs were transfected into HEK293T cells. Five days after transfection, targeted deep sequencing revealed C to T base editing ranging from 2.5% to 44.0% (Figure 4A). We also replaced the nickase form of SpCas9 with SlugCas9n in the ABEmax ^28^ plasmid to generate TadA-TadA*-SlugCas9n (SlugABEmax) (Figure 4B). SlugABEmax and the respective gRNAs were transfected into HEK293T cells. Five days after transfection, targeted deep sequencing revealed that A to G base editing ranging from 2.3% to 41.5% (Figure 4B). Collectively, these data demonstrated that SlugCas9 can be used for base editing at varying efficiencies.

**Figure 4.**
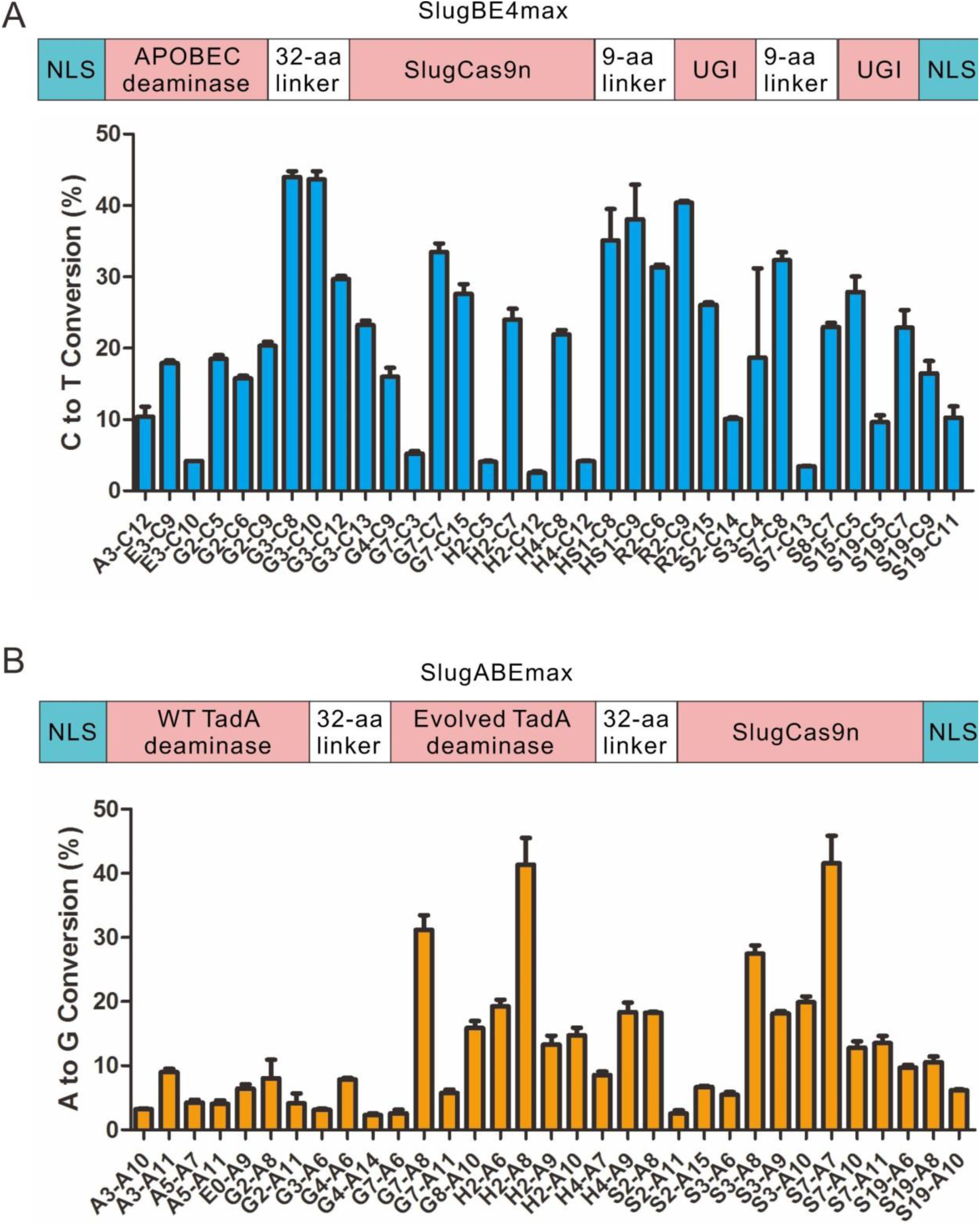
Base editing with SlugCas9. (A) SlugBE4max induces C to T conversions for 34 human genomic loci. Schematic of the SlugBE4max is shown above. n=2. (B) SlugABEmax induces A to G conversions for 33 human genomic loci. Schematic of the SlugABEmax is shown above. n=2.

### Engineering of SlugCas9 for improved specificity

Next, we evaluated the specificity of SlugCas9 by using a previously developed GFP-activation approach^18^. We first generated a panel of gRNAs with dinucleotide mutations (Figure 5A) along the protospacer to detect the specificity of SlugCas9. SlugCas9 showed robust activity for both on-target and off-target cleavage (Figure 5A), indicating that it has strong off-target effects. Recently, Tan et al. introduced quadruple mutations into SaCas9 to generate a highly specific SaCas9 variant (SaCas9-HF) ^29^. To improve the specificity of SlugCas9, we used pairwise alignment and introduced the corresponding quadruple mutations (R247A, N415A, T421A and R656A) into SlugCas9 to generate SlugCas9-HF (Figure 5B). Interestingly, SlugCas9-HF dramatically improved specificity (Figure 5A).

**Figure 5.**
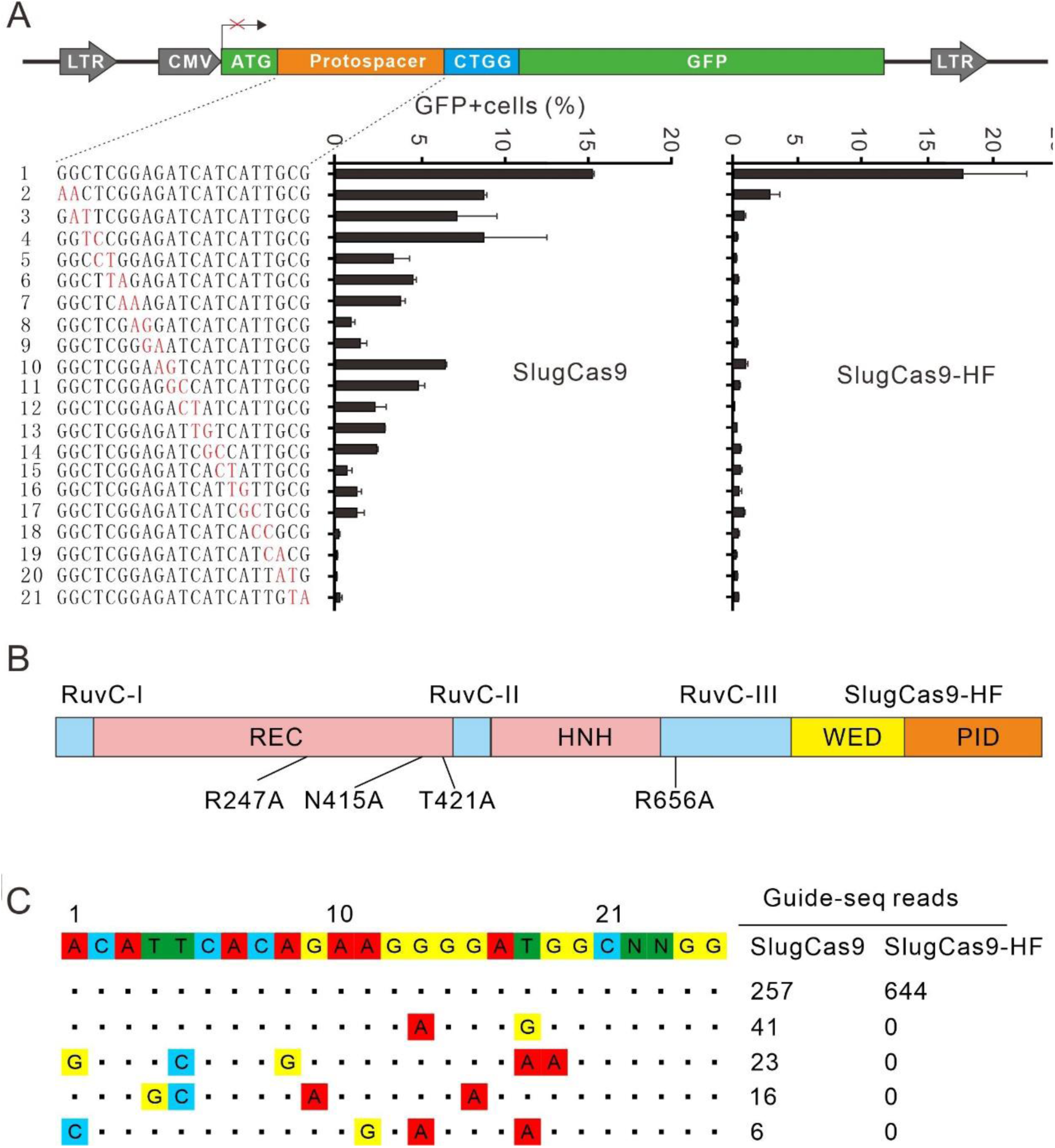
Analysis of SlugCas9 and SlugCas9-HF specificity. (A) Schematic of the GFP-activation assay for specificity analysis is shown on the top. A panel of gRNAs with dinucleotide mutations is shown below. Each gRNA activity for SlugCas9 and SlugCas9-HF is analyzed based on GFP expression. n≥2. (B) Schematic of the SlugCas9-HF. Mutations are shown below. (C) Off-targets for EMX1 locus are analyzed by GUIDE-seq. Read numbers for on- and off-targets are shown on the right. Mismatches compared with the on-target site are shown and highlighted in color.

To compare genome-wide off-target effects of SlugCas9 and SlugCas9-HF, GUIDE-seq was performed ^30^. Following transfection of SlugCas9 variants, the gRNA plasmid, and the GUIDE-seq oligos, we prepared libraries for deep sequencing. Sequencing and analysis revealed that on-target cleavage occurred for both Cas9 nucleases, reflected by the high GUIDE-seq read counts (Figure 5C). Four off-target sites were identified for SlugCas9 but no off-target sites were identified for SlugCas9-HF with this particular protospacer. In summary, these data indicated that the specificity of SlugCas9-HF was significantly improved.

To ensure no other effects were the cause of increased specificity, we compared the SlugCas9-HF activity side-by-side to SlugCas9. The editing results revealed that SlugCas9 and SlugCas9-HF displayed similar efficiency although efficiencies varied for specific loci (Figure 3A-B). These data demonstrated that SlugCas9-HF improved specificity without compromising activity.

### Engineering of SlugCas9 for expanding targeting scope

A previous study has shown that a SaCas9 variant (SaCas9-KKH) with triple mutations (E782K/N968K/R1015H) relieved the PAM sequence requirement at position 3 of the 6 base pair PAM ^19^. SlugCas9 already contained a K967 residue, which correlates to position N968 of SaCas9. To broaden the targeting scope of SlugCas9, we introduced two mutations (Q782K/R1014H) that correspond to E782K and R1015H on SaCas9, resulting in SlugCas9-KH. We performed a PAM screening assay to reveal that SlugCas9-KH preferred an NNRG (R=A or G) PAM (Figure 6A). Next, we introduced the corresponding triple mutations (Q783K/Y968K/R1015H) into ShaCas9, resulting in ShaCas9-KKH. The PAM screening assay also revealed that ShaCas9-KKH had relieved PAM preference and now preferred an NNRRC PAM (Figure 6B). Next, we introduced triple mutations (Q782K/Y966K/R1013H) into SlutrCas9, resulting in SlutrCas9-KKH. Interestingly, SlutrCas9-KKH relieved PAM restriction at position 3. The PAM screening assay revealed that the SlutrCas9-KKH now preferred an NNNRR PAM (Figure 6C). Since all these modified Cas9 nucleases are restricted by two or three nucleotides at PAM, we did not further test their activity.

**Figure 6.**
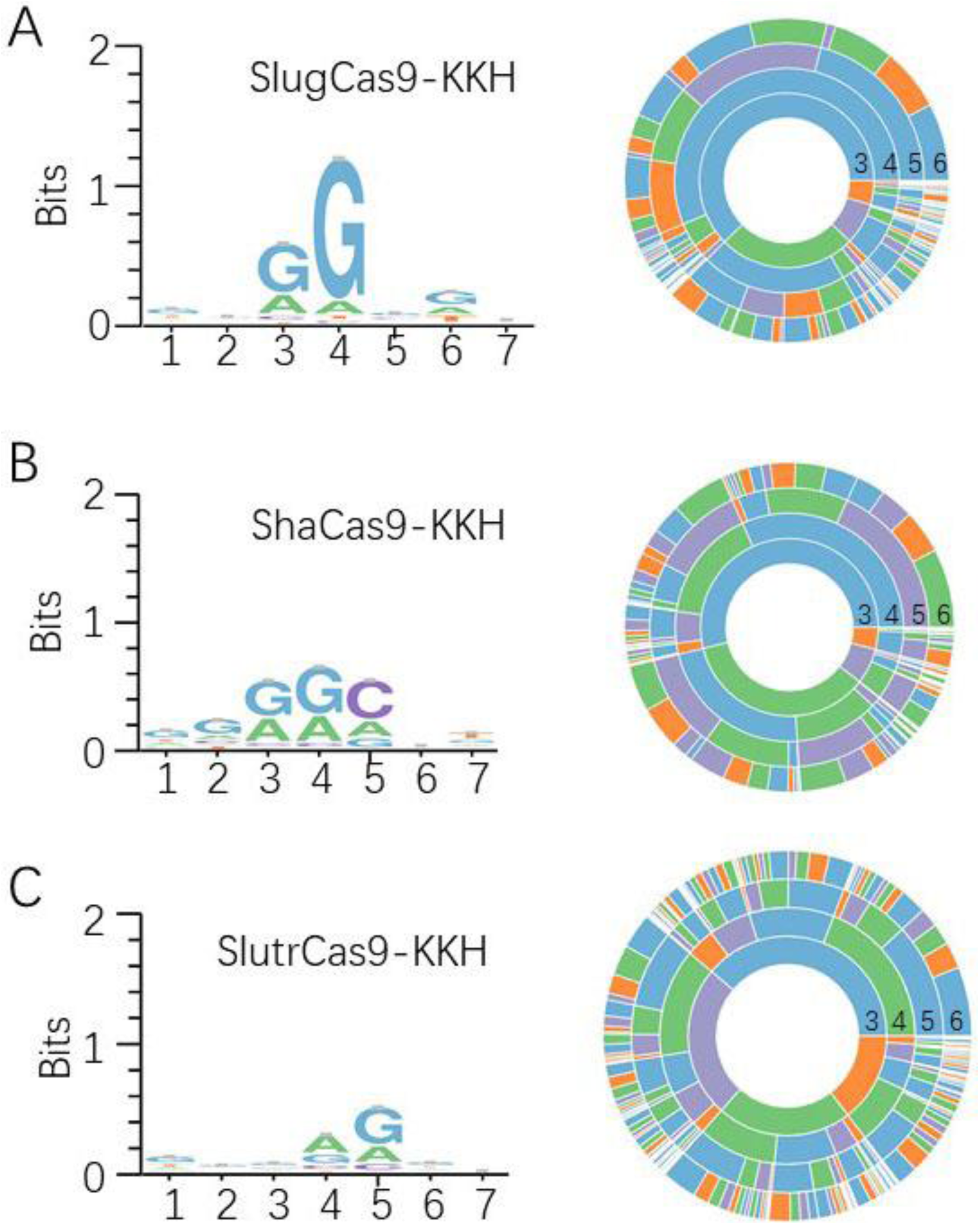
PAM analysis of engineered SaCas9 orthologs. (A) WebLogo and PAM wheel for SlugCas9-KH. (B) WebLogo and PAM wheel for ShaCas9-KKH. (C) WebLogo and PAM wheel for SlutrCas9-KKH.

### A chimeric Cas9 nuclease for specific and efficient genome editing

We previously replaced the SaCas9 PAM interacting domain (PID) with the *Staphylococcus Auricularis* Cas9 (SauriCas9) PID to generate a chimeric Cas9 nuclease (Sa-SauriCas9), which displayed high fidelity and broad targeting scope ^18^. In this study, we used the same approach to generate a chimeric Sa-SlugCas9 (Figure 7A). The GFP –activation PAM screening assay revealed that the Sa-SlugCas9 preferred an NNGG PAM (Figure 7B-C). The GFP-activation assay also revealed that the Sa-SlugCas9 specificity was similar to SaCas9 (Figure 7D). We tested the activity of Sa-SlugCas9 at a panel of endogenous loci in HEK293T cells. Overall, Sa-SlugCas9 activity was comparable to SaCas9 and SlugCas9 (Figure 7E), and was more efficient than Sa-SauriCas9.

**Figure 7.**
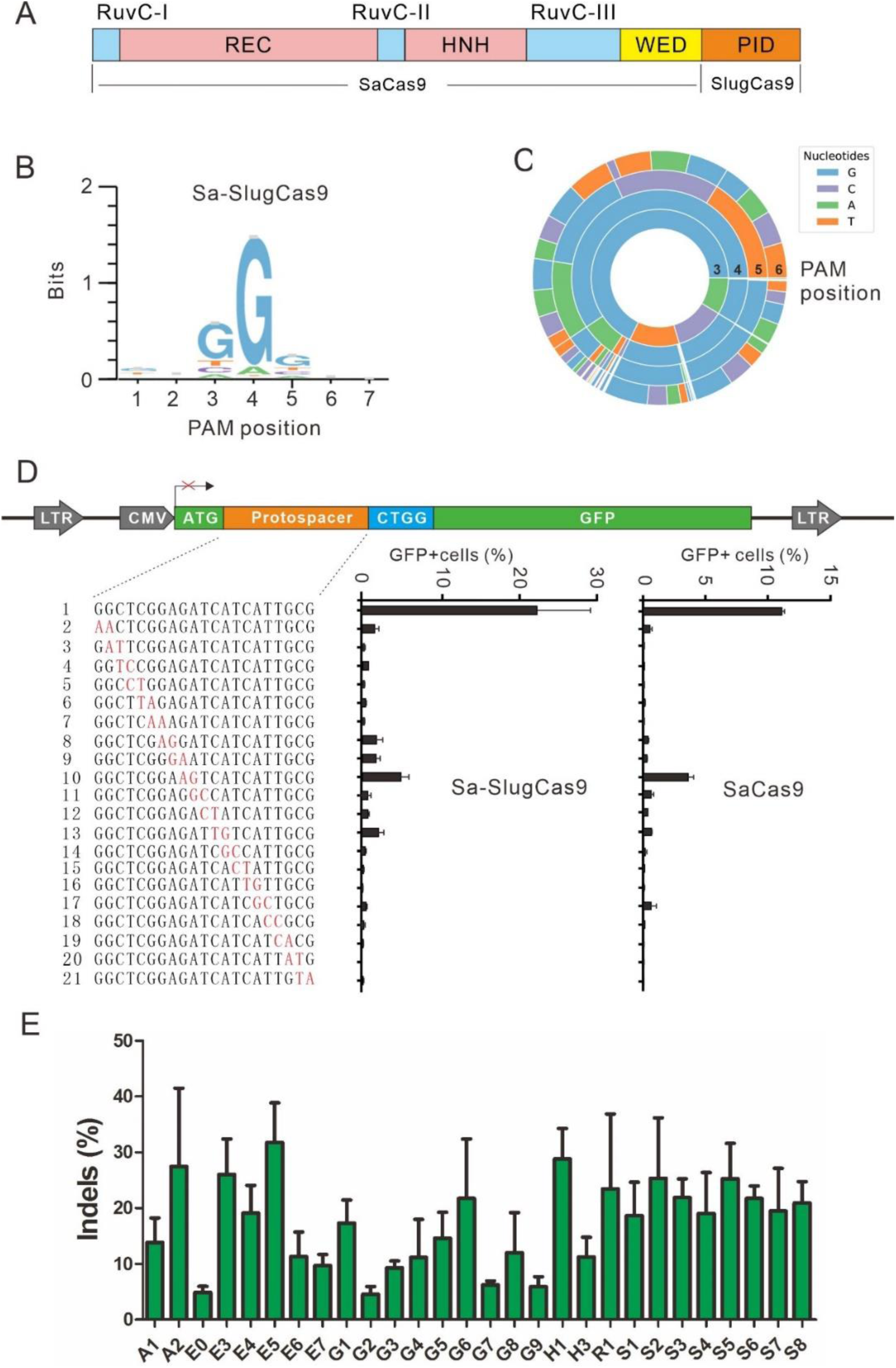
Characterization of Sa-SlugCas9 for genome editing. (A) Schematic diagram of Sa-SlugCas9. (B-C) WebLogo and PAM wheel of Sa-SlugCas9 are generated from deep-sequencing data. (D) Specificity of Sa-SlugCas9 and SaCas9 is measured by the GFP-activation assay. A panel of gRNAs with dinucleotide mismatches (red) is shown below. (E) Sa-SlugCas9 generates indels for a panel of 28 endogenous loci (n=2).

## Discussion

Small Cas9 nucleases can be packaged into the size-limited AAV vector and hold great promise for gene therapy. Although several small Cas9 nucleases have been developed for genome editing, each of them have their own limitations, such as low activity, low specificity or small targeting range ^2, 16, 18^. SaCas9 is the first small Cas9 ortholog that has been delivered by an AAV vector for *in vivo* genome editing ^2^. SaCas9 displayed higher activity amongst other small Cas9 nucleases, but it has limited utility due to its long PAM sequence requirement (NNGRRT). Engineered SaCas9 variants have increased the targeting scope (recognizing NNNRRT PAM) ^19, 31^, but this increase in targeting scope often comes at the cost of reduced on-target activity. Three type II-C Cas9 orthologs have been used for mammalian genome editing, including N. meningitidis Cas9 (NmeCas9) ^14, 15^, Campylobacter jejuni Cas9 (CjeCas9) ^16^ and N. meningitidis Cas9 (Nme2Cas9) ^17^. However, these three Cas9 nucleases generally display low editing efficiency ^32, 33^. In addition, NmeCas9 and CjeCas9 require longer PAMs, N4GAYW/N4GYTT/ N4GTCT and N4RYAC, respectively.

We recently developed SauriCas9 for mammalian genome editing ^18^. SauriCas9 recognizes a simple NNGG PAM and displays similar activity to SaCas9. However, the specificity of SauriCas9 is lower than that of SaCas9. We also generated a chimeric Sa-SauriCas9 nuclease which recognizes the same simple NNGG PAM and displays similar specificity to SaCas9, but its editing activity is low. In this study, we identified additional three SaCas9 orthologs for genome editing – SlugCas9, ShaCas9, and SlutrCas9. Interestingly, SlugCas9 recognizes a simple NNGG PAM and displays similar activity to SaCas9. By replacing the SaCas9 PID with SlugCas9 PID, we developed Sa-SlugCas9, which recognizes the simple NNGG PAM and displays similar specificity to SaCas9. Importantly, the activity of Sa-SlugCas9 is similar to SaCas9 and is significantly higher than Sa-SauriCas9. In addition, we also developed a high-fidelity version of SlugCas9, SlugCas9-HF, which recognizes the simple NNGG PAM and displays high specificity and activity. We anticipate that these novel genome editing tools will be important for both basic research and clinical applications due to its increased activity, minimal PAM requirement, high on-target:off-target ratio, and its small size for delivery efforts.

## Materials and methods

### Cell culture and transfection

HEK293T, HeLa and Human foreskin fibroblast (HFF) cells were maintained in Dulbecco’s Modified Eagle Medium (DMEM) supplemented with 10% FBS (Gibco). N2a cells were maintained in 45% DMEM and 45% Opti-MEM supplemented with 10% FBS (Gibco). HCT116 cells were maintained in McCoy’s 5A supplemented with 10% FBS (Gibco). A375 cells were maintained in RPMI-1640 supplemented with 10% FBS (Gibco), All the cells were supplemented with 100 U/ml penicillin, and 100 mg/ml streptomycin, and cultured at 37 °C and 5% CO2. For SlugCas9, Sa-SlugCas9, ShaCas9 and SlutrCas9 PAM sequence screening, HEK293T cells were plated into 10 cm dishes and transfected at 50∼60% confluency with Cas9-gRNA-expressing plasmid (15 μg) using 30 ul of Lipofectamine 2000 (Life Technologies) in Opti-MEM. For genome editing comparisons of SaCas9, SlugCas9, SlugCas9-HF, Sa-SlugCas9, ShaCas9 and SlutrCas9, cells were seeded on 48-well plates and transfected with Cas9 plasmid (300 ng) and gRNA plasmid (200 ng) using 1 ul of Lipofectamine 2000 (Life Technologies) in Opti-MEM according to the manufacturer’s instructions. For base editing capability of SlugCas9, HEK293T cells were seeded on 48-well plates and transfected with SlugABEmax/SlugBE4max plasmid (300 ng) and gRNA plasmid (200 ng) using 1 ul of Lipofectamine 2000 (Life Technologies) in Opti-MEM(Gibco) according to the manufacturer’s instructions.

### Plasmid construction

#### Cas9 expression plasmid construction

The plasmid pX601 (addgene#61591) was amplified by primers px601-F/px601-R to remove SaCas9. The human codon-optimized Cas9 gene were synthesized by HuaGene (Shanghai, China) and cloned into the pX601 backbone by NEBuilder assembly tool (NEB). The sequence of each Cas9 was confirmed by Sanger sequencing (GENEWIZ, Suzhou, China).

#### SlugBE4max and SlugABEmax plasmid construction

The primers ABEmax-F/ABEmax-R and AncBE4max-F/ AncBE4max-R was used to amplified pCMV_ABEmax_P2A_GFP (Addgene #112101) and pCMV_AncBE4max (Addgene #112094) respectively, to removed SpCas9n. SlugCas9n was amplified from SlugCas9 plasmid by primers ABE-SlugCas9(D10A)-F/ABE-SlugCas9(D10A)-R and BE4-SlugCas9(D10A)-F/BE4-SlugCas9 (D10A)-R, respectively. The PCR products were cloned into pCMV-ABEmax_P2A_GFP and pCMV_AncBE4max backbone to obtain SlugABEmax and SlugBE4max, respectively.

#### PAM sequence analysis

Twenty base-pair sequences (AAGCCTTGTTTGCCACCATG/GTGAGCAAGGGCG AGGAGCT) flanking the target sequence (GAACGGCTCGGAGATCATCATTGCG NNNNNNN) were used to fix the target sequence. GCG and GTGAGCAAGGGCG AGGAGCT were used to fix 7-bp random sequence. Target sequences with in-frame mutations were used for PAM analysis. The 7-bp random sequence was extracted and visualized by WebLog3^34^ and PAM wheel chart to demonstrate PAMs ^11^.

#### Genome editing for endogenous sites

HEK293T cells were seeded into a 48-well plate and transfected with Cas9 plasmids (300 ng) + gRNA plasmids (200 ng) by Lipofectamine 2000 (1ul) (LifeTechnologies). Cells were collected 3-5 days after transfection. For HeLa, A375, HCT116, HFF, and N2a cells, we transfected SlugCas9 (300 ng) + gRNA plasmids (200ng) by Lipofectamine 3000 (1ul) (Life Technologies). Cells were collected 5-7 days after transfection. Genomic DNA was isolated, and the target sites were PCR-amplified by nested PCR amplification and purified by a Gel Extraction Kit (QIAGEN) for deep sequencing.

#### Base editing with SlugABEmax/SlugBE4max

HEK293T cells were seeded into 48-well at a density of 20,000 cells/250 ul and transfected with SlugABEmax/ SlugBE4max (300 ng) and gRNA plasmids (200 ng). Cells were collected and the genomic DNA was isolated 3-5 days after transfection. Genomic DNA was isolated, and the target sites were PCR amplified by nested PCR amplification and purified by a Gel Extraction Kit (QIAGEN) for deep sequencing.

#### Test of Cas9 specificity

To test the specificity of SlugCas9, SlugCas9-HF, Sa-SlugCas9, ShaCas9 and SlutrCas9, we generated a GFP-reporter cell line with NNGGA (CTGGA) PAM. The cells were seeded into 48-well and transfected with Cas9 plasmids (300ng) and gRNA plasmids (200ng) by 1 ul of Lipofectamine 2000. Three days after editing, the GFP-positive cells were analyzed on the Calibur instrument (BD). Data were analyzed using FlowJo. For SaCas9 specificity, we re-analyzed the previously generated data ^18^.

#### AAVs production

HEK293T cells were seeded at approximately 60-70% confluency in a 10cm dish the day before transfection. For each well, 4 μg of Cas9-gRNA expressing plasmid, 4 μg of pAAV-RC (Gene-Bank: AF369963), and 8 μg of pAAV-helper were transfected using 160 μl of PEI (0.1% m/v, Poly-sciences, Cat# 23966 [pH 4.5]). Media was changed 6-8 hours after transfection. After 72 hours, cells are scrapped and poured into a 15-mL conical centrifuge tube. Spin at 4,000g at 4°C for 5 minutes, and transfer supernatant into a new 15-mL tube. Resuspend the cell pellet in 1 mL of PBS Buffer. Transfer to a new 15-mL conical tube. Freeze in liquid nitrogen for 1-2 minutes, and thaw at 37°C for 3-4 minutes, and repeat 3 times. Spin at 4,000g at 4°C for 10 minutes. Mix the 2 supernatants together and filter with a 0.45-μm polyvinylidene fluoride filter. Add one-half volume of the mixed solution (1MNaCl + 10% PEG8000), and incubate at 4°C overnight. After centrifugation at 4°C for 2 hours at 12,000g, discard the flow-through, and add 500 μL of chilled PBS and add the flowthrough into a 12 well with approximately 60-80% confluency HEK293T.

#### GUIDE-seq

GUIDE-seq experiments were performed as described previously ^30^, with minor modifications. Briefly, 2×10^5^ HEK293T cells were transfected with 1 μg of SlugCas9/ SlugCas9-HF, 0.5 μg of gRNA plasmids, and 100 pmol of annealed GUIDE-seq oligonucleotides by electroporation and then seeded into 12 wells. Electroporation voltage, width, and number of pulses were 1,150 V, 30 ms, and 1 pulse, respectively. Genomic DNA was extracted with a DNeasy Blood and Tissue kit (QIAGEN) 5-7 days after transfection according to the manufacturer’s protocol. Prepare the genome library and deep sequencing.

#### Test of gRNA activity in mouse cells

Six gRNAs targeting mouse Myh6 Exon 3 were selected and each of them was cloned into BsaI-digested Sa-SlugCas9 expression vector. Each gRNA activity was tested in the N2a cells. Transfection was performed with Lipofectamine 2000 according to the manufacturer’s instructions (Thermo Fisher Scientific). In brief, N2a cells were seeded onto a 48-well plate one day before transfection, and transfection was performed at about 70%-80% confluency. Sa-SlugCas9 with gRNA plasmid (500ng) were transfected.

#### Preparing of Cas9 and gRNA mRNA

Sa-SlugCas9 and gRNA DNA templates containing a T7 promoter were obtained by PCR-amplification on Sa-SlugCas9 plasmid using NEB Q5® High-Fidelity DNA Polymerase. Purified PCR products were used as templates for in vitro transcription using HiScribe™ T7 High Yield RNA Synthesis Kit (NEB, New England Biolabs). Cas9 mRNA and gRNAs were then purified using RNAClean XP(Beckman Coulter) and resuspended in RNase-free water. Both of them were stored in −80°C.

#### Quantification and statistical analysis

All the data are shown as mean ± SD. Statistical analyses were conducted using Microsoft Excel. Two-tailed, paired Student’s t-tests were used to determine statistical significance when comparing 2 or 3 groups. A value of p < 0.05 was considered to be statistically significant. (*P < 0.05, **P < 0.01, ***P < 0.001).

## Acknowledgments

This work was supported by grants from the National Natural Science Foundation of China (81870199, 81630087), the Foundation for Innovative Research Group of the National Natural Science Foundation of China (31521003), the State Key Laboratory Opening Program SKLGE1809, 111 project (B13016) and 17JC1400900.

## References

1. Wu, Z., Yang, H. & Colosi, P. Effect of genome size on AAV vector packaging. Molecular therapy: the journal of the American Society of Gene Therapy 18, 80–86 (2010).

2. Ran, F.A. et al. In vivo genome editing using Staphylococcus aureus Cas9. Nature 520, 186–191 (2015).

3. Jo, D.H. et al. Long-Term Effects of In Vivo Genome Editing in the Mouse Retina Using Campylobacter jejuni Cas9 Expressed via Adeno-Associated Virus. Molecular therapy: the journal of the American Society of Gene Therapy 27, 130–136 (2019).

4. Koo, T. et al. Functional Rescue of Dystrophin Deficiency in Mice Caused by Frameshift Mutations Using Campylobacter jejuni Cas9. Molecular therapy: the journal of the American Society of Gene Therapy 26, 1529–1538 (2018).

5. Jinek, M. et al. A programmable dual-RNA-guided DNA endonuclease in adaptive bacterial immunity. Science 337, 816–821 (2012).

6. Cong, L. et al. Multiplex genome engineering using CRISPR/Cas systems. Science 339, 819–823 (2013).

7. Mali, P. et al. RNA-guided human genome engineering via Cas9. Science 339, 823–826 (2013).

8. Xie, Y. et al. An episomal vector-based CRISPR/Cas9 system for highly efficient gene knockout in human pluripotent stem cells. Scientific reports 7, 2320 (2017).

9. Wang, D. et al. Optimized CRISPR guide RNA design for two high-fidelity Cas9 variants by deep learning. Nature communications 10, 4284 (2019).

10. Wang, B. et al. krCRISPR: an easy and efficient strategy for generating conditional knockout of essential genes in cells. Journal of biological engineering 13, 35 (2019).

11. Leenay, R.T. et al. Identifying and Visualizing Functional PAM Diversity across CRISPR-Cas Systems. Molecular cell 62, 137–147 (2016).

12. Chatterjee, P., Jakimo, N. & Jacobson, J.M. Minimal PAM specificity of a highly similar SpCas9 ortholog. Science advances 4, eaau0766 (2018).

13. Chatterjee, P. et al. A Cas9 with PAM recognition for adenine dinucleotides. Nature communications 11, 2474 (2020).

14. Esvelt, K.M. et al. Orthogonal Cas9 proteins for RNA-guided gene regulation and editing. Nature methods 10, 1116–1121 (2013).

15. Hou, Z. et al. Efficient genome engineering in human pluripotent stem cells using Cas9 from Neisseria meningitidis. Proceedings of the National Academy of Sciences of the United States of America 110, 15644–15649 (2013).

16. Kim, E. et al. In vivo genome editing with a small Cas9 orthologue derived from Campylobacter jejuni. Nature communications 8, 14500 (2017).

17. Edraki, A. et al. A Compact, High-Accuracy Cas9 with a Dinucleotide PAM for In Vivo Genome Editing. Molecular cell 73, 714–726 e714 (2019).

18. Hu, Z. et al. A compact Cas9 ortholog from Staphylococcus Auricularis (SauriCas9) expands the DNA targeting scope. PLoS biology 18, e3000686 (2020).

19. Kleinstiver, B.P. et al. Broadening the targeting range of Staphylococcus aureus CRISPR-Cas9 by modifying PAM recognition. Nature biotechnology 33, 1293–1298 (2015).

20. The UniProt, C. UniProt: the universal protein knowledgebase. Nucleic acids research 45, D158–D169 (2017).

21. Nishimasu, H. et al. Crystal Structure of Staphylococcus aureus Cas9. Cell 162, 1113–1126 (2015).

22. Dang, Y. et al. Optimizing sgRNA structure to improve CRISPR-Cas9 knockout efficiency. Genome biology 16, 280 (2015).

23. Gaudelli, N.M. et al. Programmable base editing of A*T to G*C in genomic DNA without DNA cleavage. Nature 551, 464–471 (2017).

24. Komor, A.C., Kim, Y.B., Packer, M.S., Zuris, J.A. & Liu, D.R. Programmable editing of a target base in genomic DNA without double-stranded DNA cleavage. Nature 533, 420–424 (2016).

25. Thuronyi, B.W. et al. Continuous evolution of base editors with expanded target compatibility and improved activity. Nature biotechnology 37, 1070–1079 (2019).

26. Richter, M.F. et al. Phage-assisted evolution of an adenine base editor with improved Cas domain compatibility and activity. Nature biotechnology 38, 883–891 (2020).

27. Kim, Y.B. et al. Increasing the genome-targeting scope and precision of base editing with engineered Cas9-cytidine deaminase fusions. Nature biotechnology 35, 371–376 (2017).

28. Koblan, L.W. et al. Improving cytidine and adenine base editors by expression optimization and ancestral reconstruction. Nature biotechnology 36, 843–846 (2018).

29. Tan, Y. et al. Rationally engineered Staphylococcus aureus Cas9 nucleases with high genome-wide specificity. Proceedings of the National Academy of Sciences of the United States of America 116, 20969–20976 (2019).

30. Tsai, S.Q. et al. GUIDE-seq enables genome-wide profiling of off-target cleavage by CRISPR-Cas nucleases. Nature biotechnology 33, 187–197 (2015).

31. Ma, D. et al. Engineer chimeric Cas9 to expand PAM recognition based on evolutionary information. Nature communications 10, 560 (2019).

32. Tsui, T.K.M., Hand, T.H., Duboy, E.C. & Li, H. The Impact of DNA Topology and Guide Length on Target Selection by a Cytosine-Specific Cas9. ACS synthetic biology 6, 1103–1113 (2017).

33. Wang, Y. et al. Systematic evaluation of CRISPR-Cas systems reveals design principles for genome editing in human cells. Genome biology 19, 62 (2018).

34. Crooks, G.E., Hon, G., Chandonia, J.M. & Brenner, S.E. WebLogo: A sequence logo generator. Genome Res 14, 1188–1190 (2004).

